# Intermittent access cocaine self-administration produces psychomotor sensitization: effects of withdrawal, sex and cross-sensitization

**DOI:** 10.1101/859520

**Authors:** Crystal C. Carr, Carrie R. Ferrario, Terry E. Robinson

## Abstract

The psychomotor activating effects of drugs such as cocaine or amphetamine can change in very different ways – showing sensitization or tolerance – depending on whether they are administered more or less intermittently. This behavioral plasticity is thought to reflect, at least in part, changes in dopamine (DA) neurotransmission, and therefore, may provide insights into how repeated drug use promotes the development of substance use disorders. Indeed, the most widely used preclinical model of cocaine addiction, which involves Long Access (LgA) self-administration procedures, is reported to produce tolerance to cocaine’s psychomotor activating effects and effects on DA activity. This is cited as evidence in support of the view that in addiction, drug-seeking and-taking is motivated to overcome this DA deficiency and associated anhedonia. In contrast, Intermittent Access (IntA) cocaine self-administration is more effective than LgA in producing addiction-like behavior, but sensitizes DA neurotransmission. There is, however, very little information concerning the effects of IntA experience on the psychomotor activating effects of cocaine. The purpose of the studies reported here, therefore, was to determine whether IntA experience produces psychomotor sensitization with similar characteristics to that produced by the intermittent, noncontingent administration of cocaine. It did. The psychomotor sensitization produced by IntA experience with cocaine: (1) was greater after a long (30 days) vs short (1 day) period of withdrawal; (2) was greater in females than males; and (3) resulted in cross-sensitization to another psychomotor stimulant drug, amphetamine. This pattern of cocaine experience-dependent plasticity favors an incentive-sensitization view of addiction.

## Introduction

Many drugs of abuse have psychomotor activating effects, indicated by an increase in locomotor activity and rearing at low doses, and at higher doses by focused stereotyped behaviors, such as repetitive head, limb and oral movements-effects thought to be largely due to their ability to increase dopamine (DA) neurotransmission in the dorsal and ventral striatum (Wise and Bozarth 1987, for review). Historically, interest in the psychomotor activating effects of drugs stems in part because such effects provide an indicator of DA activity, and in part because it has been argued that, *“the reinforcing effects of drugs, and thus their addiction liability, can be predicted from their ability to induce psychomotor activation”* (Wise and Bozarth 1987; p.474). Furthermore, when many drugs of abuse are administered repeatedly and intermittently there is a progressive increase (sensitization) in their psychomotor activating effects, which is associated with increased DA neurotransmission (for reviews see, Robinson and Becker 1986; Kalivas and Stewart 1991; Stewart and Badiani 1993; Vezina 2004). Such sensitization-related forms of neurobehavioral plasticity have been hypothesized to contribute to addiction (Robinson and Berridge 1993).

Many early studies of the phenomenon of psychomotor sensitization involved noncontingent drug injections given by an experimenter, and it became important, therefore, to determine if sensitization-related changes in brain and behavior are also induced when drugs are self-administered. Indeed, there are many reports that drug (here we focus on cocaine) self-administration not only induces psychomotor sensitization (Hooks et al. 1994; Phillips and Di Ciano 1996; Zapata et al. 2003) but also sensitization to its motivational effects (Deroche et al. 1999; Vezina et al. 2002; Morgan et al. 2006; Lack et al. 2008) and sensitization of DA neurotransmission, especially when animals are tested after a period of withdrawal (Hooks et al. 1994; Zapata et al. 2003; Wiskerke et al. 2016). However, most of these early studies of sensitization produced by cocaine self-administration experience involved relatively limited or Short Access (ShA) conditions (although see Ferrario et al., 2005), and over the years it became increasingly apparent that such conditions are not especially effective in producing addiction-like behavior (for review see, Ahmed 2011). This led to considerable effort to develop more realistic preclinical models of addiction (Ahmed and Koob 1998; Deroche-Gamonet et al. 2004) that can be uniquely valuable for isolating persistent drug-induced changes in brain and behavior that may promote addiction, which is difficult to do in humans.

Currently, the most widely used rodent model of cocaine addiction utilizes Long-Access (LgA) intravenous (IV) self-administration procedures, introduced by Ahmed and Koob in 1998. This involves allowing rats to self-administer cocaine for 6 hrs or more in a single daily session. Relative to rats tested under more limited, or Short Access (ShA, 1-2 h/day) conditions, LgA experience is reported to produce a number of addiction-like behaviors, including escalation in drug-intake (Ahmed and Koob, 1998), high motivation for drug (Paterson and Markou 2003; Morgan et al. 2006; Wee et al. 2008; Ben-Shahar et al. 2008), continued drug-seeking in the face of an adverse consequence (Vanderschuren and Everitt 2004; Pelloux et al. 2007), and a high propensity to relapse (Mantsch et al. 2004; Kippin et al. 2006). LgA is also reported to reduce evoked DA release in the ventral striatum (Calipari et al. 2013, 2014a), and decrease the ability of cocaine to induce striatal DA overflow and to inhibit DA uptake via the DAT (Ferris et al. 2011; Calipari et al. 2013, 2014a), all of which is accompanied by a decrease in cocaine’s psychomotor activating effects – the opposite of sensitization (Calipari et al. 2013, 2014a). Thus, studies using LgA procedures have been cited in support of the view that in addiction, brain reward systems are rendered *hypo*sensitive because of blunted DA neurotransmission. By this view drug-seeking and -taking are primarily motivated by a desire to overcome a deficiency in DA, and associated anhedonia (e.g., Koob and Volkow 2016; Volkow et al. 2016).

In contrast, studies using a more recently developed Intermittent Access (IntA) cocaine self-administration procedure paint a very different picture (Zimmer et al. 2012; for reviews, Allain et al. 2015; Kawa et al. 2019a). IntA involves the use of successive drug available periods, interspersed with periods when drug is not available, which produces a series of ‘spikes’ in brain cocaine concentrations within a session. This is thought to better reflect the temporal pattern of use in humans, especially during the transition to addiction (Ward et al. 1997; Beveridge et al. 2012). Importantly, IntA experience not only produces the addiction-like behaviors described above, but in many instances is *more* effective in doing so than LgA, despite much less total drug consumption (Kawa et al. 2016; Allain et al. 2017; Allain and Samaha 2018; Allain et al. 2018; James et al. 2019; Kawa et al. 2019b). Furthermore, IntA experience sensitizes striatal DA neurotransmission (Calipari et al. 2013, 2014b, 2015; Kawa et al. 2019b). Thus, studies using IntA procedures are more consistent with an incentive-sensitization view of addiction, which posits that, “addiction is caused primarily by drug-induced sensitization in the brain mesocorticolimbic systems [including DA] that attribute incentive salience to reward-associated stimuli” (Robinson and Berridge 2008, p. 3137). By this view, drug-seeking and taking in addiction is primarily motivated by a *hyper*-responsive DA system and sensitized “wanting” (Robinson and Berridge 1993).

The question addressed here is whether IntA experience also produces psychomotor sensitization. A few studies suggest yes, based on measures of locomotor activity within-subjects during IntA sessions (Allain et al. 2017; Allain and Samaha 2018; Algallal et al. 2019), but this phenomenon has not been well characterized. We asked, therefore, whether the psychomotor sensitization produced by IntA experience has similar characteristics to that described in many studies utilizing experimenter-administered drugs. Specifically, we asked whether, as with experimenter-administered drug, the psychomotor sensitization produced by IntA experience is expressed to a greater extent after long vs short periods of withdrawal (Robinson and Camp 1987; Paulson et al. 1991; Paulson and Robinson 1995; Grimm et al. 2001); whether females show greater sensitization than males (Glick and Hinds 1984; Robinson 1984; Van Haaren and Meyer 1991; for review, Becker et al. 2006); and whether treatment with one psychostimulant, cocaine, produces cross-sensitization to another, amphetamine (Akimoto et al. 1990; Schenk et al. 1991; Hirabayashi et al. 1991; Shanks et al. 2015).

## General Materials and Methods

### Subjects

Male and female Sprague-Dawley rats (Envigo, Haslett, MI), ~55 days old upon arrival, were housed individually in a climate-controlled colony room on a reverse 12-h light/12-h dark cycle. All training and testing were conducted during the dark period. Rats were given 1 week to acclimate to the colony room before any manipulation. Water and food were available *ad libitum* until 2 days before the first self-administration session. At this time, all rats were mildly food restricted to maintain a stable body weight. This was done to prevent excessive weight gain, which is unhealthy, particularly in adult male rats (Rowland 2007). All procedures were approved by the University of Michigan Institutional Animal Care and Use Committee.

### Intravenous catheter surgery

Intravenous (IV) catheters were surgically implanted as described previously (Crombag et al., 2000). Briefly, rats were anesthetized with ketamine hydrochloride (90 mg/kg, IP) and xylazine (10 mg/kg, IP), and an indwelling catheter was secured into the jugular vein. The catheter exited via a port situated just above the shoulder blades. All rats were given antisedan (1 mg/kg, IP) following completion of the surgery, to rapidly reverse the effects of xylazine. The analgesic carprofen was given at the start of the surgery and again the two days following surgery (5mg/kg, SC). In addition, catheters were flushed daily with 0.2 ml sterile saline containing 5 mg/ml gentamicin sulfate (Vedco, MO) for the duration of the experiment. Catheter patency was tested prior to behavioral testing, and periodically throughout an experiment (e.g., if responding for cocaine suddenly declined), by administering an IV infusion of 0.1 mL of methohexital sodium (10 mg/ml in sterile water, JHP Pharmaceuticals). If a rat did not become ataxic within 10 seconds of the infusion it was removed from the study.

### Cocaine self-administration

All self-administration testing took place in Med Associates chambers (22 × 18 × 13 cm; St Albans, VT, USA) located within sound-attenuating cabinets. Each chamber was equipped with a ventilating fan, which masked background noise, a tone generator, and a red house light. For all studies, active responses resulted in an IV infusion of cocaine hydrochloride (NIDA, 0.4 mg/kg/infusion, weight of the salt, in 50 *μ*l of sterile saline delivered over 2.6-sec), whereas inactive responses were recorded but had no consequence. In addition, each infusion was accompanied by a compound tone and light stimulus, which continued throughout the time out period (see individual experimental design below for additional details). For all studies, infusions were given on a fixed ratio 1 schedule of reinforcement, and one session was conducted per day.

### Acquisition of cocaine self-administration

To ensure that all rats received the same amount of drug and cue exposures during initial training, an infusion criteria (IC) procedure was used, as described previously (Saunders and Robinson 2010). During each IC session rats were allowed to take a fixed number of infusions (e.g., 10 infusions = IC10), with the number of infusions available increasing across days as each rat met the infusion criteria (e.g., IC10, IC20, IC40). During initial acquisition the time out period was 20-sec (including infusion time). Rats were removed from the chambers when the criterion was met, and thus session length varied from animal to animal.

### Intermittent and Long Access cocaine self-administration

Intermittent access (IntA) procedures were similar to those previously described (Zimmer et al. 2012; Kawa et al. 2016). Briefly, each session consisted of alternating Drug-Available and No Drug-Available periods lasting 5-min and 25-min, respectively. The house light was on at the start of the session, and the first Drug-Available period was signaled by the house light turning off. During Long Access (LgA) sessions, drug was available throughout the entire 6-h session. For both IntA and LgA the time out period was 7.6-sec. Importantly, each IntA session consisted of 12 Drug-Available and 12 No Drug-Available periods, resulting in a 6-h session length, and therefore, both LgA and IntA groups were exposed to the self-administration chambers for the same amount of time each day.

### Assessing psychomotor activity

Depending on the experiment, psychomotor activity was assessed following experimenter-administered and/or self-administered cocaine. In all experiments involving experimenter-administered cocaine, behavior was recorded in test chambers with walls made of black, expanded PVC (68.58 × 33.02 × 66.04 cm), and grey wire mesh grid floors. Each chamber was equipped with a camera (CVC-130R, Speco Technologies, Amityville, NY, USA) suspended ~18 cm above the center of each test chamber. To assess the behavioral response to self-administered cocaine *during* an IntA session, rats were allowed to administer three infusions of cocaine during each Drug-Available period within the self-administration chamber. In these studies, each self-administration chamber was equipped with a camera mounted on the back wall of the sound-attenuating chamber. Additional details regarding groups, habituation, and doses tested are given within each experiment below.

Cocaine and amphetamine both produce psychomotor activation, which can manifest as increases in locomotor activity, as well as increases in repetitive stereotyped behaviors (e.g., head movements), during which time locomotion may decrease (Lyon and Robbins 1975). Locomotor activity in the psychomotor test chambers was quantified using TopScan motion-tracking software (High-Throughput Option Version 3.00, Clever Sys Inc., Reston, Virginia, USA) or EthoVision XT software (Version 11.5, Noldus Information Technology b.v., Wageningen, The Netherlands). For these automated analyses (TopScan and EthoVision) user-defined areas were located at the far left and far right side of the chamber, and *crossovers* from one area to the other counted using center-point detection, as an estimate of locomotor activity. Importantly, direct comparisons were only made between measures made using the same software and approach.

Locomotor activity in the self-administration chambers was scored by visual examination of videos. For this, the experimenter was blind to experimental conditions and the number of infusions available was capped to three so that all rats took the same amount of cocaine (see also Exp 2 below). Behavior was then quantified during each following 25-min No Drug-Available period. The chamber was divided into two equal halves and locomotor activity was determined by again counting the number of crossovers (center-point) from one side of the test chamber to the other, in 5-min bins.

Stereotyped behavior was examined using two different measures. One, behavior was rated using a scale adapted from Ellinwood and Balster, 1974 and Robinson et al., 1988 (see Table 1). Two, a parametric measure of the intensity of stereotypy was obtained by quantifying the frequency of head movements during periods when the rat was ‘in place’, as described by Ferrario et al. (2005). Periods ‘in place’ were defined as periods when both back paws remained in the same place for a minimum of 2-sec. The frequency of head movements was then determined by dividing the total number of head movements by the total time spent in place. For both rating and frequency measures, one 30-sec sample of behavior was assessed every 5-min during the first 40-min following each cocaine injection, resulting in eight 30-sec samples/dose/rat.

**Table 1.**
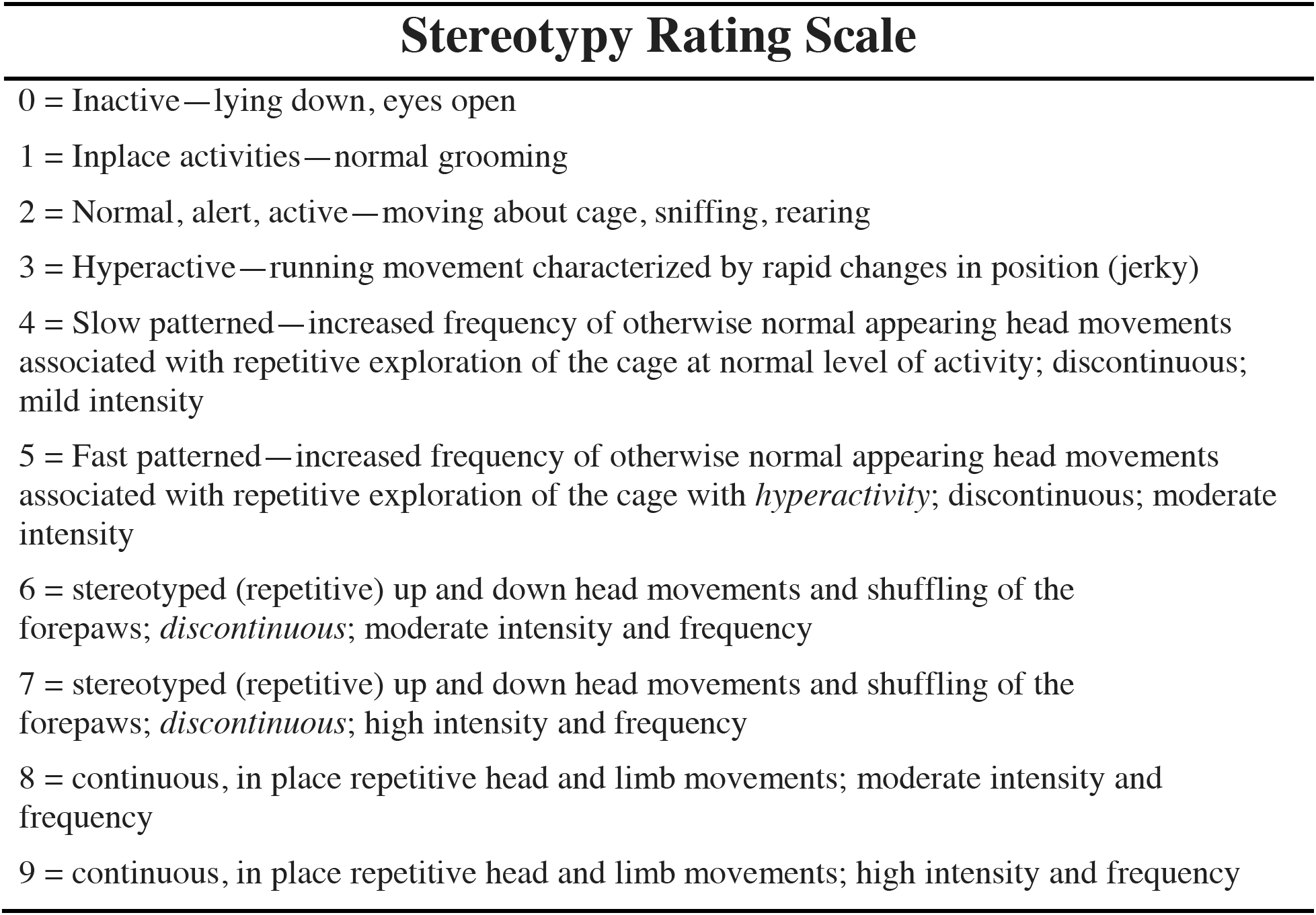
Rating scale used to assess the psychomotor activating effects of cocaine. Adapted from Ellinwood & Balster, 1974 and Robinson et al., 1988.

### Experiment 1. Does IntA and LgA cocaine self-administration experience produce psychomotor sensitization that varies as a function of withdrawal period?

The psychomotor response to experimenter-administered (IP) cocaine was assessed in male rats prior to any self-administration experience (Baseline) and then again on challenge test days conducted after 1 and 30 days of withdrawal (WD1, WD30) from IntA or LgA cocaine self-administration experience (see Timeline shown in Fig. 1a).

**Fig. 1.**
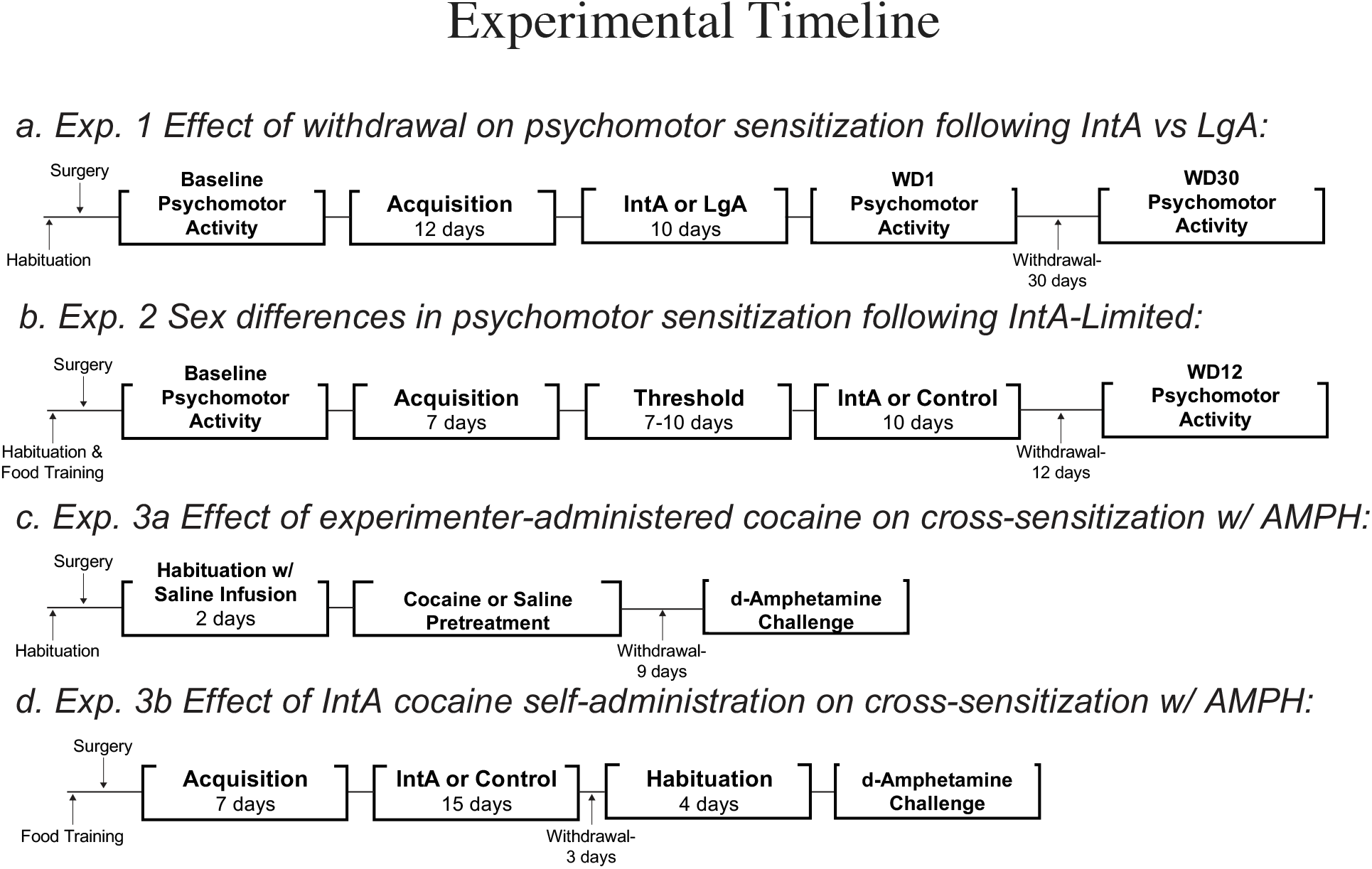
Flow diagrams for each experiment. Exp, Experiment; IntA, Intermittent Access; LgA, Long Access; Amph, *d*-Amphetamine.

Rats were first habituated to the psychomotor activity test chambers by placing them into the chambers for one hour on two consecutive days. Next, rats underwent IV catheter surgery followed by 7 days of recovery as described in General Methods above. Baseline cocaine-induced psychomotor activity was then assessed as follows. Rats were placed into the psychomotor test chambers and left undisturbed for 30-min. They were then removed and given an IP injection of saline, and placed back into the chamber for 30-min. This was followed by an IP injection of 7.5 mg/kg of cocaine, then 1-h later 15 mg/kg, and after an additional 1.5-h a 30 mg/kg injection, as described previously (Ferrario et al. 2005; Oginsky et al. 2016). Behavior was video-recorded throughout and testing was conducted in red light conditions. Next, rats underwent acquisition for cocaine self-administration using the IC procedure described above. For this study, the self-administration chambers were equipped with one active and one inactive nose-poke port (right/left counter-balanced). Active responses resulted in an infusion of cocaine, the illumination of the active nose-poke port and presentation of a tone. The tone and nose-poke light remained on during the infusion and the 20-sec time out period. Each rat was given 2 sessions of IC10, 3 sessions of IC20, and 6 sessions of IC40. Rats were then assigned to IntA (n=9) or LgA (n=10) groups, counterbalanced by average session duration during the last three IC40 sessions.

### Experiment 2: Are there sex differences in the psychomotor sensitization produced by IntA cocaine self-administration experience?

Repeated, intermittent treatment with experimenter-administered psychomotor stimulant drugs produces greater psychomotor sensitization in female than male rats (Glick and Hinds 1984; Robinson 1984; Van Haaren and Meyer 1991; for review, Becker et al. 2006). The purpose of this experiment was to determine if this is also the case following IntA cocaine self-administration experience (Fig. 1b).

For this experiment, the self-administration chambers were equipped with an active and an inactive retractable lever, rather than nose poke ports, in part to prevent responding during No Drug-Available periods, which could interfere with the quantification of drug-induced psychomotor activity (see below). In addition, because females often acquire cocaine self-administration more readily than males, rats in this study first underwent food self-administration training, in an attempt to reduce differences in acquisition between the sexes. For food self-administration training (30-min per session, 2-5 sessions total), a food cup was located in the center of the front wall of the chamber. Each response on the active lever resulted in the delivery of one food pellet (45 mg, banana flavored pellets; BioServe, #F0059, Frenchtown, NJ, USA), paired with illumination of the cue light above the active lever. Responses on the inactive lever had no consequence and active/inactive levers were right/left counterbalanced. During this time, rats were also habituated to the psychomotor test chambers (1-h/day, 2 consecutive days). After food training, rats underwent IV catheter surgery followed by recovery and baseline IV psychomotor testing as described below.

All rats underwent two days of habituation to the psychomotor test chambers and experimenter administered drug injection procedure. On these days the rats were first placed in the chambers for 30-min; they were then removed and placed into a square plastic holding cage. Their IV catheter was then attached to a length of PE20 tubing connected to a 1.0 ml syringe mounted on a Harvard Apparatus syringe pump (Holliston, MA). The syringe pump was then used to infuse 60 *μ*l of saline over 5-sec (142 *μ*l/min). The tubing was then detached from the catheter and animals were quickly placed back into the psychomotor test chambers for an additional 30-min. Three additional saline infusions were administered separated by 30-min, for a total of 4 saline infusions. On the baseline test day, rats were again placed in the chamber for 30-min followed by one saline infusion. Next, rats were given three infusions of increasing doses of cocaine (0.25, 0.5. and 1.0 mg/kg, IV), with 30-min between each infusion. The infusion itself consisted of 30*μ*l of saline, followed by 10*μ*l of cocaine, and another 20*μ*l of saline for a total infusion volume of 60*μ*l. This ensured that the infusion volume was larger than the dead space of the IV catheter. Behavior was video-recorded throughout.

Following baseline psychomotor testing, rats underwent acquisition of cocaine self-administration using the IC procedure. Because they had already undergone food training, IC training was reduced to 2 sessions of IC10 followed by 5 sessions of IC20. Each infusion was accompanied by the illumination of a cue light located above the active lever. The cue light remained illuminated during the 20-sec time out period. In addition, the active lever was retracted during each time out in order to familiarize the rat with lever retraction.

Female rats have been reported to self-administer more cocaine, and are more motivated to do so, relative to males (Roberts et al. 1989; Lynch and Taylor 2004; Roth and Carroll 2004; Lynch et al. 2005; Cummings et al. 2011; Smith et al. 2011). Therefore, we next sought to determine if this were the case under our test conditions. To this end, rats were next trained to self-administer using a within-session threshold procedure, as described previously (Oleson et al. 2011; Bentzley et al. 2013). At the start of these 110-min sessions (signaled by the house light being off and insertion of the levers, which remained extended throughout the entire session), rats earned infusions on an FR-1 schedule and the dose of cocaine available decreased every 10-min on a quarter logarithmic scale (1.28, 0.72, 0.40, 0.23, 0.13, 0.072, 0.040, 0.023, 0.013, 0.007, 0.004 mg/kg/infusion). Dose was manipulated by changing the duration of the infusion, and the cue light remained illuminated during the duration of each infusion (7-10 session). There was no signaled time out period, but additional infusions could not be earned while an infusion was being given.

Next, rats were assigned to IntA (males n=10, females n=9) or Control groups (males n=10, females n=9). Rats in the Control group did not receive any additional self-administration experience, but each day they were removed from their home cage and placed in holding chambers for the same amount of time as rats in the IntA group were in the self-administration chambers. In order to ensure that there were no differences in the total amount of cocaine consumed between males and females in the IntA group (which could itself produce differences in psychomotor sensitization), we used a modified IntA procedure similar to that described previously (Allain et al. 2018). Briefly, rats were limited to three infusions per 5-min Drug-Available period (IntA-Limited). The cue light was illuminated for the duration of each infusion (2.6-sec). The No Drug-Available period started as soon as the allotted infusions were self-administered, or after 5-min, whichever came first, and was signaled by illumination of the house light and retraction of both levers. Each IntA-Limited session consisted of 8 Drug-Available and 8 No Drug-Available periods. Locomotor activity in response to self-administered cocaine was evaluated during IntA-Limited sessions 1, 3, 7 and 10, within subjects. Finally, the psychomotor response to an experimenter-administered IV infusion of cocaine was evaluated 12 days after IntA-Limited testing, using a between-subjects design comparing IntA-Limited and Control groups. The same multi-dose procedure described for initial baseline testing was used (see also General Methods for details of psychomotor measures).

### Experiment 3: Does IV cocaine produce cross-sensitization to IV amphetamine?

Experimenter-administered cocaine given IP induces cross-sensitization to IP amphetamines (Akimoto et al. 1990; Hirabayashi et al. 1991; Shanks et al. 2015), but similar cross-sensitization between IV cocaine and amphetamine had not been established. Therefore, we first determined whether experimenter-administered *IV* infusions of cocaine would also produce cross-sensitization to subsequent IV amphetamine in females (Exp. 3a; see also Timeline Fig 1c). We then asked whether IntA cocaine self-administration experience would as well (Exp. 3b; see also Timeline Fig 1d).

### Exp. 3a. Amphetamine cross-sensitization to experimenter-administered IV cocaine

Female rats were first habituated to the psychomotor activity test chambers (1-h per day, 2 days), and then underwent IV catheter surgery as described above. They were next habituated to the experimenter-administered IV infusion procedure (2 days, 30-min followed by a single IV infusion of saline; 60 *μ*l over 5-sec) and returned to their home cage after an additional 30-min. Rats were then assigned to a Saline-(n=10) or a Cocaine-treated (n=11) group. On each treatment day, rats were placed into the chambers for 30-min, and then given an IV infusion of cocaine (1 mg/kg) or saline (60 *μ*l delivered over 5-sec). Behavior post infusion was video recorded for 30-min before returning rats to their home cages. Infusions were given every other day for a total of 7 infusions. Rats then experienced 9 days of withdrawal, during which time catheters were flushed daily with 0.2ml of saline containing 5 mg/ml gentamicin sulfate. On WD10 the response to IV challenge infusions of *d*-amphetamine was examined.

On the amphetamine challenge day, rats were placed in the chamber for 30-min and then all rats (both Saline and Cocaine pretreated groups) were given an infusion of saline. After 30-min, all rats were given an IV infusion of 0.3 mg/kg of *d*-amphetamine. 2.5-h later rats received an infusion of 0.6 mg/kg of *d*-amphetamine (infusion volume: 60 *μ*l delivered over 5-sec). Behavior was video-recorded throughout testing and recordings ended 3.5-h after the last infusion (see also General Methods).

### Exp. 3b. Amphetamine cross-sensitization following IntA cocaine self-administration

Here we asked whether IntA cocaine self-administration experience produces cross-sensitization to IV amphetamine in male rats (see timeline Fig 1d). Rats were given food training followed by IV catheter surgery and recovery as described above for Exp. 2. They then underwent IC10 (2 sessions) and IC20 (5 sessions) cocaine self-administration training. Rats were randomly assigned to IntA (n=12) and Control (n=13) groups. The 15 IntA self-administration sessions were identical to those described in Exp. 2, except that rats could take an unlimited number of infusions during each of eight 5-min Drug-vailable periods. Importantly, training was staggered such that IntA and Control groups both received their amphetamine challenge test 7 days after the last cocaine self-administration session (WD7). Habituation to the psychomotor testing and infusion procedure was conducted during WD4-7, as described for Exp 3a. The amphetamine challenge test was conducted similar to that described in Exp 3a, except that doses of 0.25 mg/kg and 0.5 mg/kg *d*-amphetamine were used (infusion volume: 60 *μ*l delivered over 5-sec).

### Statistics

Prism 8.0 (GraphPad Software) was used for all parametric statistical analyses. Differences across time, session/day, or dose in locomotor activity, percent time spent in place, frequency of head movements, and infusions were all analyzed using a two-way repeated-measures analyses of variance (ANOVA) based on the general linear model. The Geisser-Greenhouse correction was applied to all factors with three or more levels (such as time and dose) to mitigate violations of sphericity. Post-hoc multiple comparisons (and Sidak corrections) were used as appropriate. Group differences in cumulative cocaine intake was analyzed using an unpaired *t*-test. Wilcox Signed Ranks Test was used to analyze the non-parametric psychomotor rating scale data (SPSS v. 24).

## Results

### Experiment 1: Does IntA and/or LgA cocaine self-administration experience produce psychomotor sensitization that varies as a function of withdrawal period?

After being trained to self-administer cocaine using an infusion criteria (IC) procedure, rats were assigned to IntA or LgA groups matched for initial performance during acquisition. There were, therefore, no group differences in the acquisition of cocaine self-administration (data not shown). As expected, during the 10 day IntA or LgA self-administration period rats in the LgA group consumed about twice as much cocaine as rats in the IntA group (Fig. 2; *t*(1,17)=7.265, *p*<0.001).

**Fig. 2.**
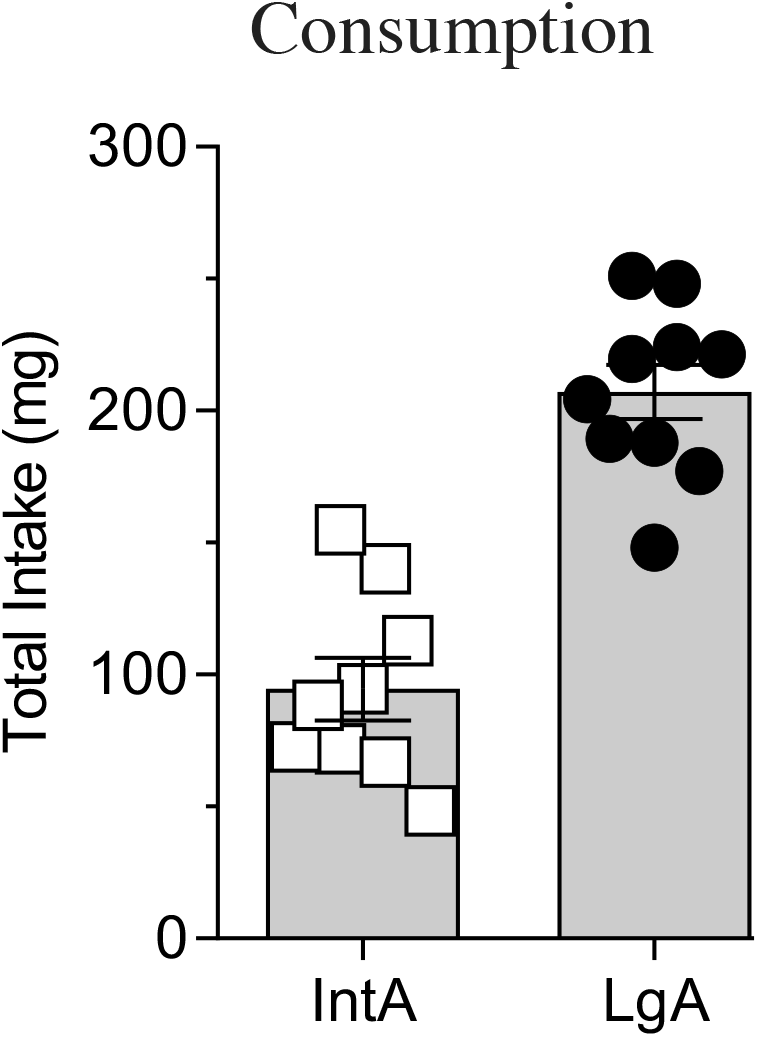
Mean (± SEMs) total cocaine intake during 10 days of access to IntA or LgA. Circles and squares represent individual rats. Rats in the LgA group (n=10) rats consumed about twice as much cocaine than those in the IntA group (n=9).

The effects of LgA or IntA cocaine self-administration experience on psychomotor activity was first assessed 1 day after the last self-administration session (WD1). The psychomotor activating effect of increasing dose of IP cocaine were compared to that seen during the Baseline test, conducted prior to IntA or LgA experience (Fig. 3). Panels a-d in Fig. 3 show the time course of locomotor activity (i.e., crossovers). Only rats with IntA experience showed a sensitized locomotor response on WD1 relative to baseline. To simplify data presentation and analysis these data were collapsed across time (crossovers/min) and expressed as dose-effect functions (Panels e, f). Compared to Baseline, IntA rats showed an enhanced (sensitized) locomotor response on WD1 across all doses tested (Fig. 3e; main effect of test day, *F*(1,8)= 11.47, *p*=0.0096; main effect of dose, *F*(2.383, 19.07)= 10.36, *p*=0.0006). In contrast, in the LgA group cocaine-induced locomotor activity increased as a function of dose (*F*(2.193, 19.74)= 7.989, *p*=0.0023), but there was no difference in locomotor activity between Baseline and WD1 (Fig. 3f; effect of test day; *F*(1,9)= 1.855, *p*=0.2063; dose X test day interaction; *F*(1.860, 16.74)= 0.8166, *p*=0.4507).

**Fig. 3.**
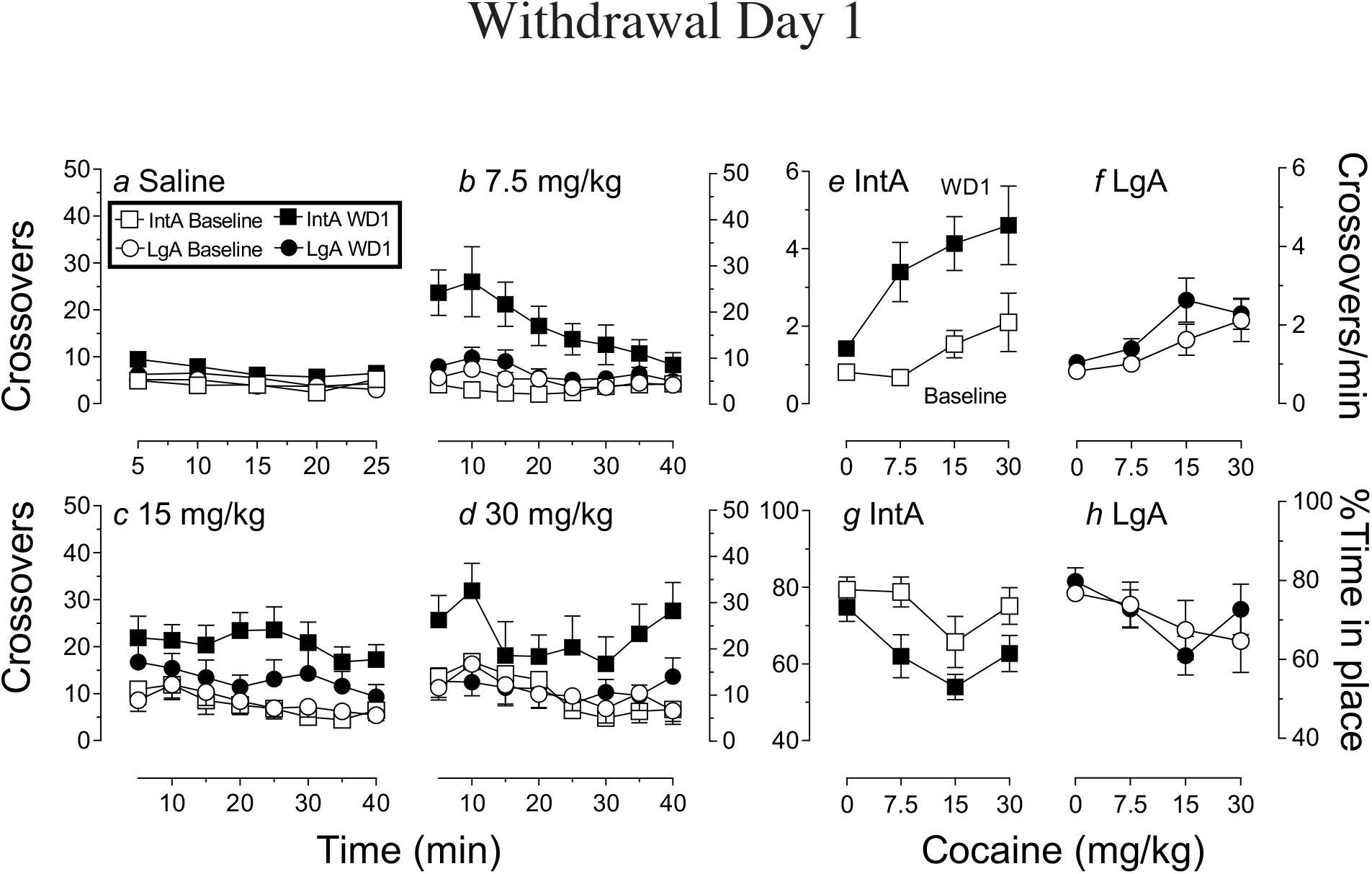
Behavioral effects of cocaine at Baseline (prior to any self-administration experience) and 1 day after the discontinuation of IntA or LgA cocaine self-administration experience (WD1). Panels a-d show the time course of locomotor activity, as assessed by crossovers. Cocaine produced greater locomotor activity following IntA, but not LgA experience. Panels e & f show these same data collapsed across time to generate a dose-effect function, which clearly demonstrates that IntA, but not LgA experience produced psychomotor sensitization (see Results for all statistics). Panels g & h show the time animals spend ‘in place’ (i.e, not locomoting) as a function of group and dose. Consistent with the measure of locomotion, cocaine produced a greater dose-dependent decrease in time in place following IntA experience, relative to baseline, but there was no effect of session in the LgA group. All data represented as mean ± SEM.

A second, non-automated measure of behavior, was derived from direct analysis of the video records. The percent of time spent in place during each test session (i.e., *not* locomoting) was calculated (Panels g and h). In the IntA group there was a dose-dependent decrease in time spent in place on both test sessions, that was greater on WD1 than at Baseline (Fig. 3g; effect of dose, *F*(2.140, 17.12)= 4.598, *p*=0.0234; effect of test day, *F*(1,8)= 10.65, *p*=0.0115), consistent with the sensitization indicated by an increase in crossovers captured by the automated measures. In the LgA group there was also a dose-dependent decrease in the time spent in place (Fig. 3h; effect of dose, *F*(2.406, 21.65)= 3.183, *p*=0.0535), but there was no difference between Baseline and WD1 (Fig. 3h; effect of test day, *F*(1,9)= 0.02, *p*=0.8899), again consistent with the automated assessment of locomotor activity.

The effect of LgA or IntA cocaine self-administration experience on psychomotor activity was next assessed on WD30 in the same rats (Fig. 4). Although at Baseline there was a dose-dependent increase in crossovers in both the IntA and LgA groups (Fig 4a-b open symbols), there was a decrease in crossovers in both groups on WD30 (Fig. 4a-b closed symbols). This was accompanied by an increase in the time spent in place (Fig. 4c-d). This pattern of behavior is inconsistent with cocaine-induced locomotor hyperactivity, but suggestive of focused stereotyped behaviors. Indeed, visual observation of videos showed animals engaged in focused stereotyped behaviors, especially at the highest dose. For this reason, measures other than locomotion was required to assess drug effects on WD30. We used two measures: 1) a rating scale to assess the psychomotor activating effects of cocaine (Table 1; adopted from Ellinwood and Balster 1974 and Robinson et al. 1988); and 2) quantification of the frequency of stereotyped head movements while otherwise ‘in place’ (Ferrario et al. 2005).

**Fig. 4.**
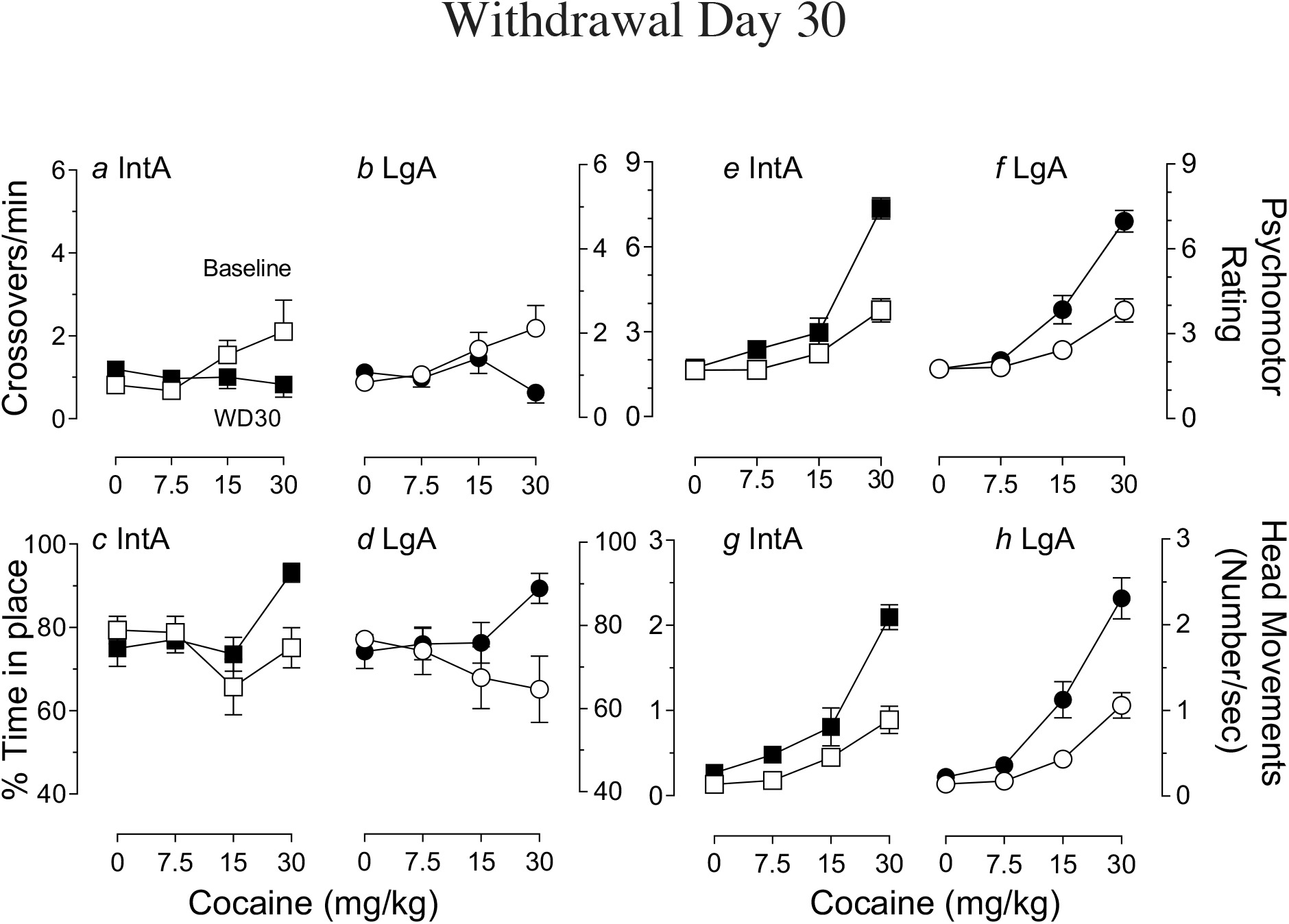
Behavioral effects of cocaine at Baseline (prior to any self-administration experience) and then 30 days after the discontinuation of IntA or LgA cocaine self-administration experience (WD30). Panels a & b show that at Baseline there was a dose-dependent increase in crossovers, but this was not evident at WD30 and in fact on WD30 there was a dose-dependent increase in time in place in both groups (Panels c & d). This pattern of behavior suggests that on WD30 cocaine produced stereotyped behaviors in both groups, which was confirmed by inspection of videos. The degree of cocaine-induced stereotypy was assess using a rating scale (Panels e & f) and by counting the frequency of stereotyped head movements (Panels g & h). Both measures showed that at this time psychomotor sensitization was evident in both the IntA and LgA groups. All data represented as mean ± SEM.

In both the IntA and LgA groups cocaine produced higher stereotypy ratings on WD30 than at Baseline (IntA: Fig. 4e; Z=−2.668, *p*=0.008; LgA: Fig. 4f; Z=−2.803, *p*=0.005). The median ratings at Baseline following 30 mg/kg were 3.75−4.0, which corresponds to hyperactivity accompanied by normal looking head movements (see Table 1). This is consistent with the locomotor hyperactivity seen at this dose (Fig. 3e, f). However, on WD30 the median ratings were 7.1-8.0, which corresponds to discontinuous to continuous stereotyped head movements of high intensity and frequency (Table 1), and is consistent with the increased time in place shown in Fig. 4c, d.

A parametric measure of stereotyped behavior was obtained by counting the frequency of head movements while a rat was otherwise ‘in place’, that is, not locomoting (Fig. 4g, h). There was a dose-dependent increase in the frequency of head movements in both groups on both test sessions, that was greater on WD30 than at Baseline (IntA: Fig. 4g; effect of dose, *F*(1.537, 12.30)= 47.95, *p*<0.0001; effect of day, *F*(1,8)=36.78, *p*=0.0003; day X dose interaction, *F*(1.735,13.88)= 7.733, *p*=0.0069; LgA: Fig. 4h; effect of dose, *F*(1.679, 15.11)= 74.41, *p*<0.0001; effect of day, *F*(1,9)= 25.26, *p*=0.0007; day X dose interaction, *F*(1.986,17.87)= 8.573, *p*=0.0025). In summary, both rating scale and head movement measures of stereotyped behavior indicated that on WD30 rats with either IntA or LgA cocaine self-administration experience expressed especially robust psychomotor sensitization (also see Ferrario et al. 2005). Thus, psychomotor sensitization was expressed on WD1 and WD30 following IntA, but only on WD30 following LgA.

### Experiment 2: Are there sex differences in the psychomotor sensitization produced by IntA cocaine self-administration experience?

There was no sex difference in days to criteria for food training (data not shown) or in the acquisition of self-administration behavior (data not shown). This is expected given that the IC procedure fixes the number of available infusions each day and rats must reach that number before moving to the next IC. During the threshold test females self-administered more cocaine than males at low doses (Fig. 5; effect of sex, *F*(1, 36)= 9.197, *p*=0.0045; effect of dose, *F*(1.762, 63.43)= 44.26, *p*<0.0001; sex X dose interaction, *F*(10, 360)= 7.673, *p*<0.0001), consistent with previous studies (Roberts et al. 1989; Cummings et al. 2011; Kawa and Robinson 2019). Therefore, in subsequent studies we used an IntA-*Limited* procedure to mitigate any effects of sex differences in total cocaine consumption on subsequent psychomotor sensitization.

**Fig. 5.**
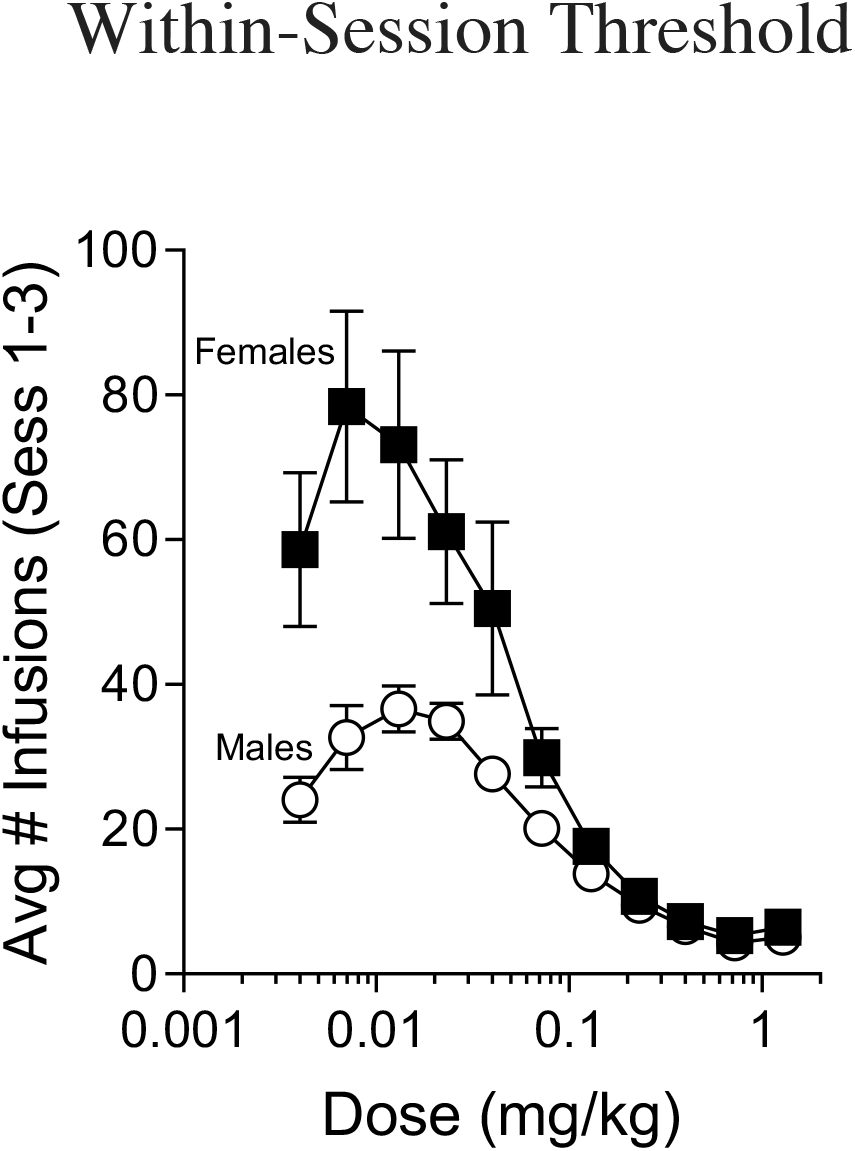
Motivation to obtain cocaine was assessed using a within-session threshold procedure after initial acquisition of cocaine self-administration. Females consumed more drug than males. All data represented as mean ± SEM.

### Psychomotor sensitization: within-subjects analysis

During the IntA-Limited cocaine-self-administration sessions the number of infusions allowed during each Drug-Available period was capped at 3, resulting in a maximum of 24 infusions per session. There were no significant group differences in the amount of cocaine consumed during IntA (data not shown; *t*(1,17)=0.9459, *p*=0.3574).

Fig. 6 shows the number of crossovers for the first three 5-min intervals during each No Drug-Available period on Sessions 1, 3, 7 and 10 of IntA-Limited testing. We examined the entire 25-min period and found that (1) group differences were confined to the first 5-10 min, and (2) activity returned to baseline within 15-25 min. Therefore, only the first 15-min are shown for the sake of clarity. There were no differences in cocaine-induced crossovers between males and females on Session 1 (Fig 6a). However, on Sessions 3, 7 and 10 self-administered cocaine produced greater psychomotor activation in females than males (Fig 6 b-d). To better display and analyze these data the number of crossovers during the first 5 min of each of the 8 No Drug-Available periods were averaged for Sessions 1, 3, 7 and 10. This captures peak locomotor activity for both males and females and is shown in Fig. 7. Both males and females showed an increase in locomotor activity across sessions (effect of session, *F*(1.758, 28.13)=11.59, *p*=0.0003). However, females showed a greater increase in locomotor activity across sessions than males, that is, greater psychomotor sensitization, as indicated by a significant sex X session interaction (*F*(3, 48)=2.895, *p*=0.0447).

**Fig. 6.**
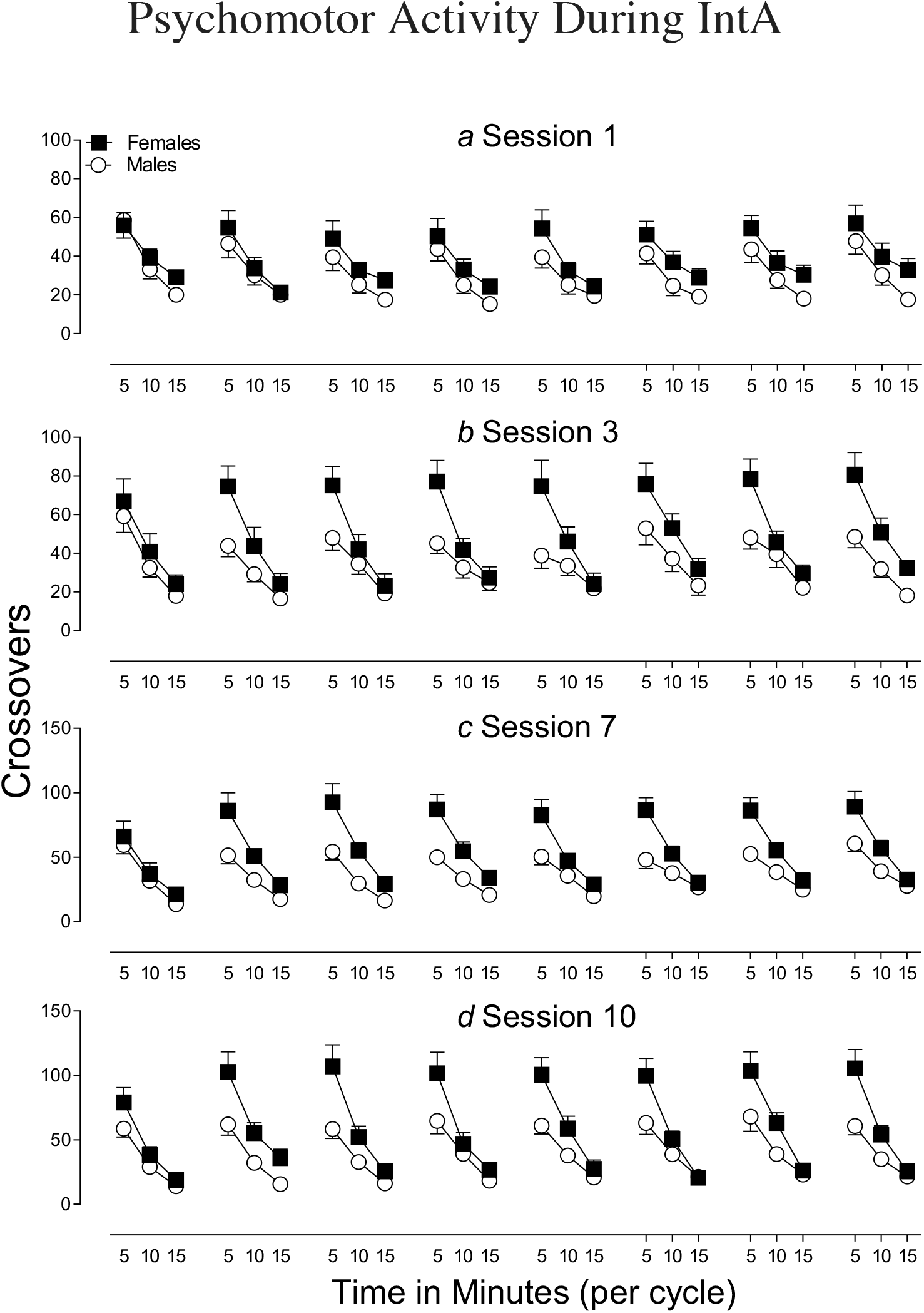
Mean (± SEM) crossovers for the first 15 min (5 min bins) after the self-administration of 3 infusions of cocaine, over 8 successive Drug-Available periods on Days 1, 3, 7 and 10 of IntA-Limited testing, in male and female rats. There was no sex difference in crossovers on Session 1 (a), but females showed greater psychomotor activation on Sessions 3, 7 and 10 (Panels, b, c & d, respectively). See Fig. 7 for analysis of the effect of Session.

**Fig. 7.**
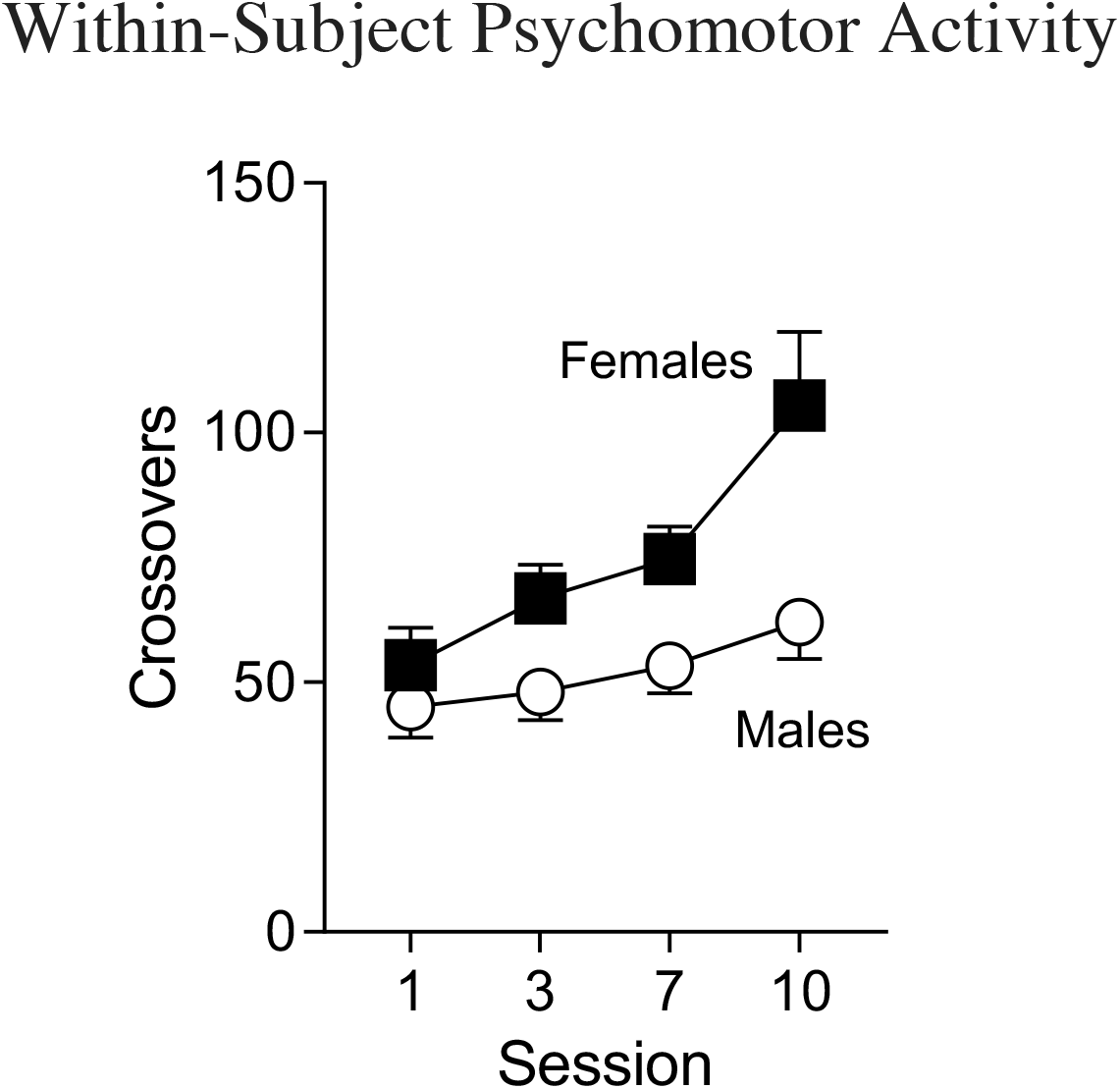
Mean (± SEM) crossovers during the first 5 min following the self-administration of 3 infusions of cocaine (averaged across the 8 daily Drug Not-Available cycles) as a function of session in male and female rats. There was an increase in crossovers as a function of session in both males and females, but the degree of psychomotor sensitization was greater in females than males.

### Psychomotor sensitization: between-subjects analysis

Psychomotor sensitization can also be assessed by comparing the magnitude of locomotor activity between subjects, providing additional corroboration of effects. For this analysis male and female rats that had IntA-Limited experience were compared to their respective Control groups. Animals in the Control groups received all the same treatments as the IntA-Limited groups, including surgery, acquisition of self-administration, and the threshold test, but no IntA-Limited experience. Twelve days after the last IntA-Limited self-administration session rats in both the Control and IntA-Limited groups received experimenter-administered IV challenge infusions of increasing doses of cocaine in the psychomotor test chambers (see Fig 1b for timeline). Note that all rats had been habituated to these test chambers, and the injection procedure, prior to acquisition of self-administration, and there were no group differences in locomotor activity at that time (data not shown). In the males, there was a dose dependent-increase in locomotor activity following IV cocaine (Fig. 8a; effect of dose, *F*(2.285, 38.84)=26.21, *p*<0.0001), but this was similar between IntA-Limited and Control groups. Females also showed a dose-dependent increase in locomotor activity (Fig. 8b; effect of dose, *F*(1.716, 27.45)=18.56, *p*<0.0001), but this was greater in the IntA than Control group (effect of group, *F*(1, 16)=10.09, *p*=0.0059; group X dose interaction, *F*(3, 48)= 2.848, *p*=0.0472). In summary, this between-subjects analysis showed that following IntA experience psychomotor sensitization was expressed in females, but not males. Taken together with within-subjects measures, these data show that IntA cocaine self-administration produces more robust psychomotor sensitization in females than males.

**Fig. 8.**
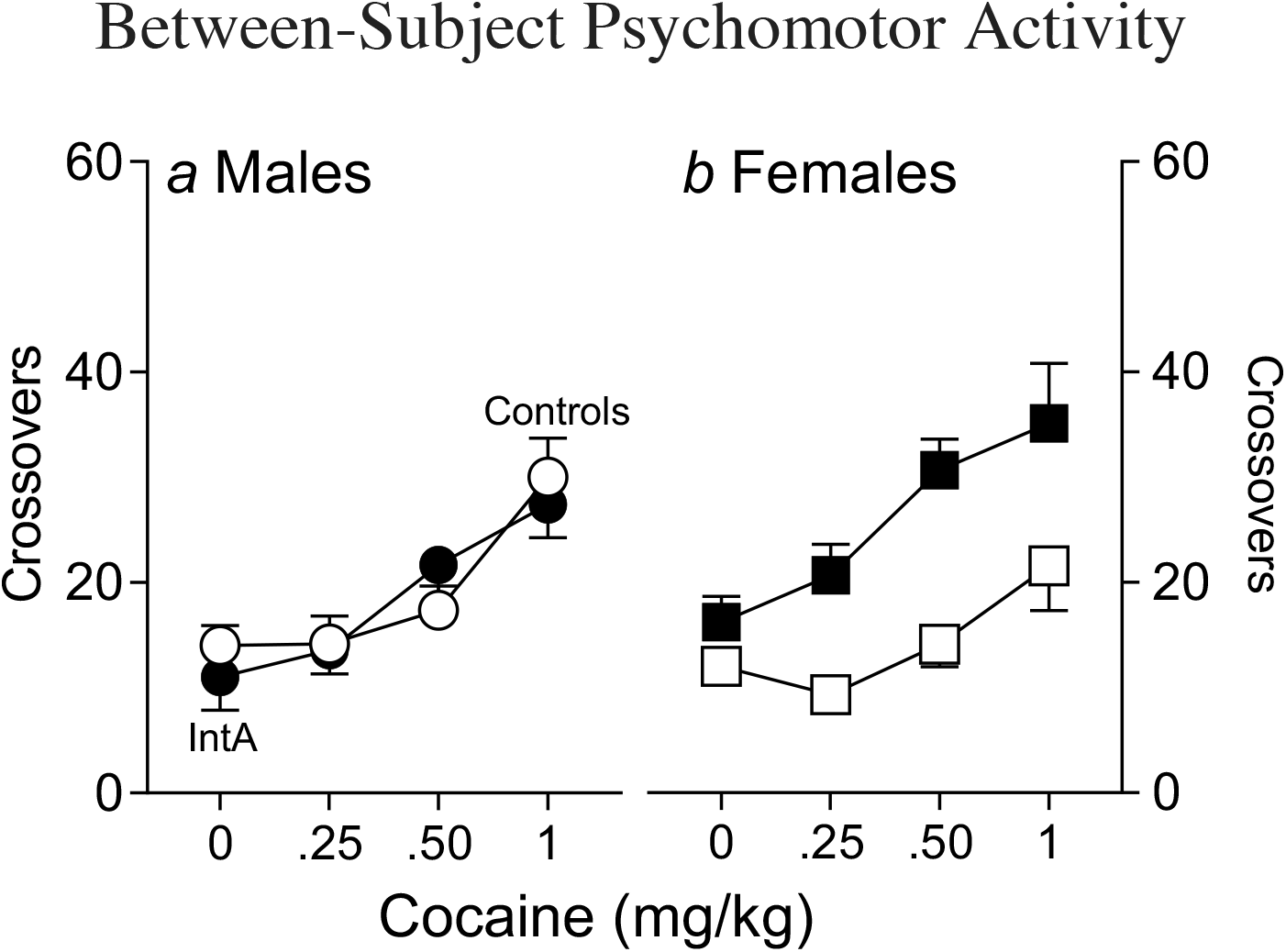
Mean (± SEM) crossovers produced by experimenter-administered IV cocaine challenge infusions in male and female rats with prior IntA cocaine self-administration experience, and Control rats that did not have IntA experience. Cocaine produced a greater dose-dependent increase in crossovers in the IntA group, relative to Controls (i.e., psychomotor sensitization), in females, but not males.

### Experiment 3: Does cocaine produce cross-sensitization to amphetamine?

Previous studies have reported that experimenter-administered cocaine produces cross-sensitization to a subsequent injection of amphetamine, but these studies all involved IP injections (Akimoto et al. 1990; Hirabayashi et al. 1991; Shanks et al. 2015). Therefore, we first determined whether experimenter-administered IV infusions of cocaine would also produce cross-sensitization (Exp 3a; Fig 1c). We then determined whether IntA cocaine self-administration experience produces cross-sensitization to amphetamine (Exp. 3b; Fig 1d).

### Experiment 3a

Cocaine produced greater psychomotor activation (crossovers) on the last (seventh) day of IV cocaine treatment than the first, indicating sensitization (Fig. 9; effect of time, *F*(3.045, 30.45)=63.15, *p*<0.0001; effect of session, *F*(1, 10)=15.6, *p*=0.0027; session X time, *F*(2.186, 21.86)=13.38, *p*=0.0001).

**Fig. 9.**
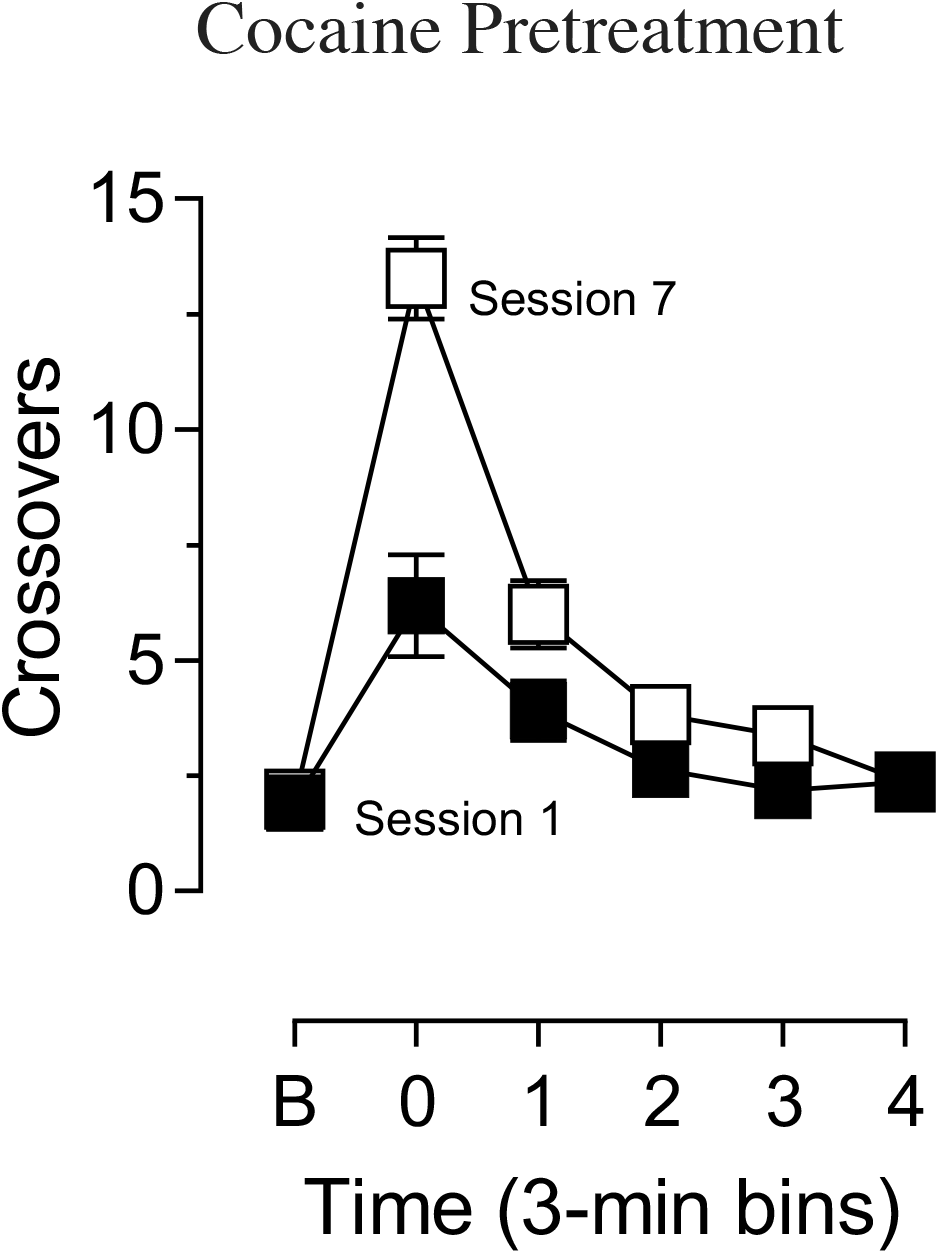
Mean (± SEM) crossovers in 3 min bins prior to (B, Baseline) and following the first and seventh infusion of experimenter-administered cocaine (1.0 mg/kg, IV; n=11). The behavioral response was greater after the seventh than the first session, indicating psychomotor sensitization.

On the challenge test day infusions of 0.3 and 0.6 mg/kg *d*-amphetamine produced greater psychomotor activation in cocaine pretreated than control groups (Fig. 10a: 0.3 mg/kg, effect of group, *F* (1, 19)=2.816, *p*=0.1097, effect of time, *F* (3.997, 75.94)=14.15, *p*<0.0001, group X time, *F* (6, 114)=3.103, *p*=0.0075; Fig. 10b: 0.6 mg/kg, effect of group, *F* (1, 19)=11.57, *p*=0.0030, effect of time, *F* (4.681, 88.93)=40.96, *p*<0.0001, group X time interaction *F* (6, 114)=2.579, *p*=0.0222). Fig. 10c shows a summary of the dose-effect relationship (effect of group, *F* (1, 19)=7.264, *p*=0.0143; effect of dose, *F* (1.729, 32.84)=42.37, *p*<0.0001; group X dose interaction, *F* (2, 38)=3.492, *p*=0.0405).

**Fig. 10.**
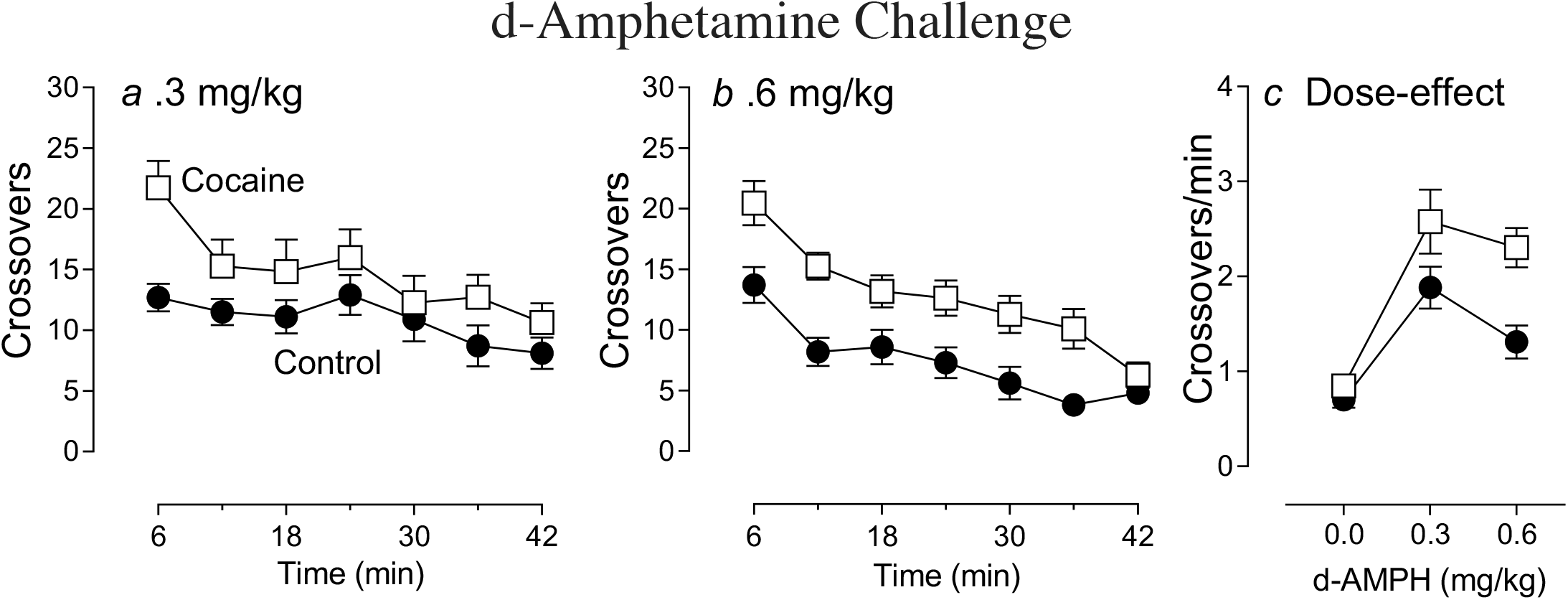
Mean (± SEM) crossovers during the amphetamine challenge test day in rats pretreated with IV experimenter-administered cocaine (open symbols) or Saline-pretreated Controls (closed symbols). Panels a & b show the time course of the locomotor response to 0.3 and 0.6 mg/kg of amphetamine IV, respectively. Panel c shows the same data averaged across each dose as a dose-effect function. Psychomotor sensitization is indicated by the greater response in the Cocaine-relative to the Saline-pretreated Control group.

### Experiment 3b

The effect of challenge infusions of *d*-amphetamine on locomotor activity (crossovers) was compared between rats that had 15 days of IntA cocaine self-administration experience compared to controls that had surgery and underwent IC self-administration training, but had no IntA experience (Fig. 1d). With increasing IntA experience rats escalated their intake during the first min of the Drug-Available periods, which was when they consumed nearly all drug (Fig. 11; effect of session, *F* (1.638, 18.02)=3.59, *p*=0.0564; effect of minute, *F* (1.233, 13.57)=192.3, *p*<0.0001; session X minute interaction (*F* (5.039, 55.43)=14.11, *p*<0.0001; post-hoc analysis revealed this was driven by minute 1). This is consistent with previous reports (Kawa et al., 2016; Kawa et al., 2018a).

**Fig. 11.**
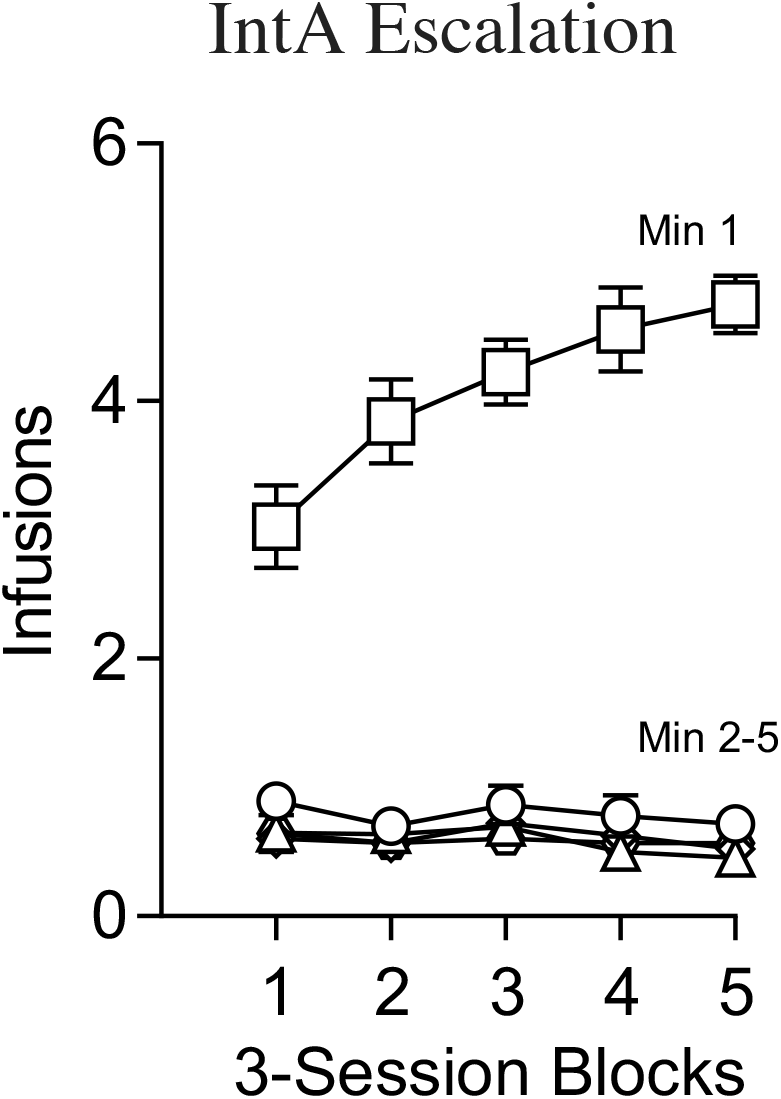
Mean (± SEM) number of infusions during each minute (Min) of the Drug-Available periods during IntA cocaine self-administration, averaged across 3 session blocks. It can be seen that (1) the rats took nearly all drug during the first minute of drug availability, and (2) that there was an increase in drug consumption with increasing IntA experience.

On the psychomotor test day the initial infusion of saline produced very little effect but the IntA group showed slightly greater activity than the control group, perhaps indicating conditioned hyperactivity (data not shown; main effect of group, *F* (1, 23)=5.118, *p*=0.0334). A challenge infusion of 0.25 mg/kg of *d*-amphetamine produced slightly more activity in the IntA group than the Controls, but this was not statistically significant (Fig 12a; effect of group, *F* (1, 23)=2.104, *p*=0.1604; group X time interaction, *F* (14, 322)=1.648, *p*=0.0655). A challenge infusion of 0.5 mg/kg of *d*-amphetamine produced more crossovers in the IntA than the Control group (Fig. 12b; Effect of group, *F* (1, 23)=8.037, *p*=0.0094; group X time interaction, *F* (14, 322)=2.339, *p*=0.0043). Fig. 12c shows the dose-effect relationship (group X dose interaction, *F* (2, 46)=4.664, *p*=0.0143). In summary, IntA cocaine self-administration experience produced cross-sensitization to *d*-amphetamine.

**Fig. 12.**
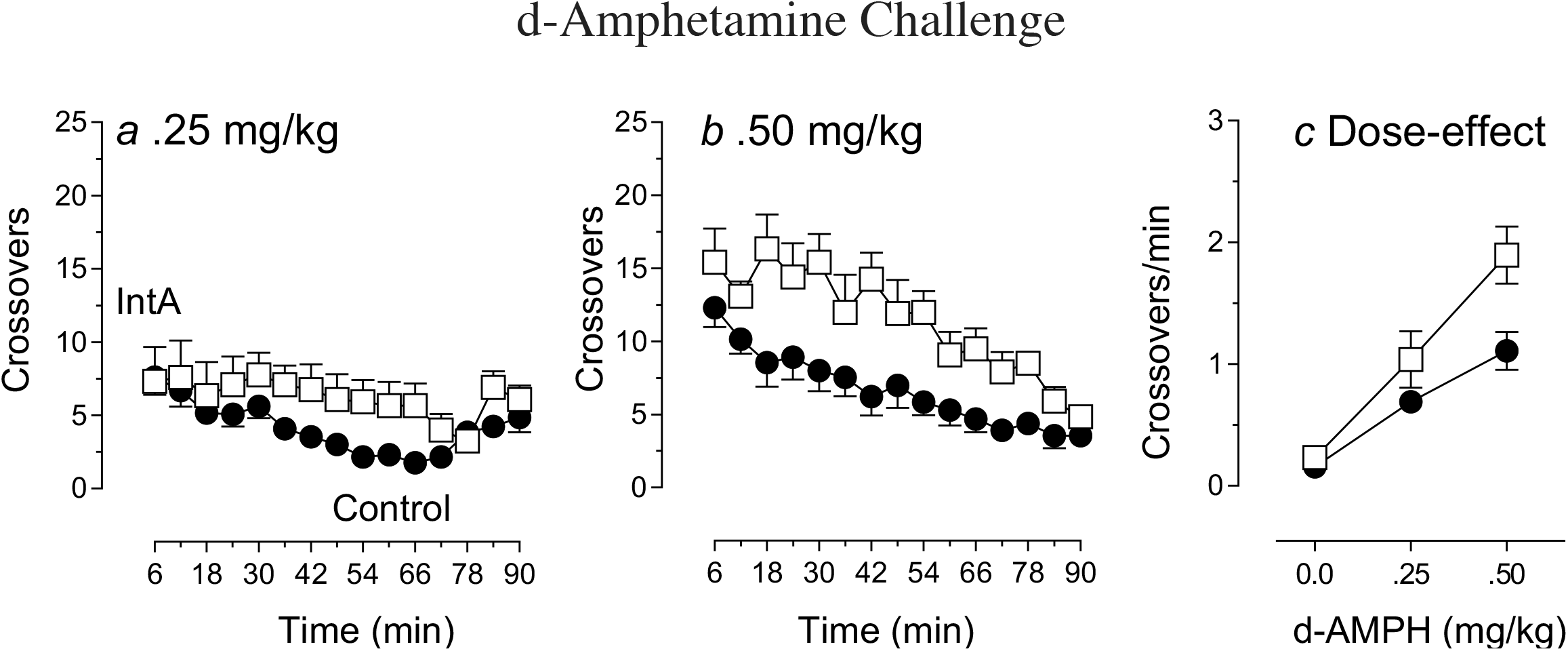
Mean (± SEM) crossovers on the *d*-amphetamine challenge test day in rats with prior IntA cocaine self-administration experience, and Control rats that did not have any IntA experience. Panels a & b show the time course of the locomotor response to 0.25 and 0.5 mg/kg of *d*-amphetamine IV, respectively. Panel c shows the same data averaged across each dose as a dose-effect function. Psychomotor sensitization is indicated by the greater response in the IntA relative to the Control group after the higher dose of *d*-amphetamine.

## Discussion

We asked whether IntA cocaine self-administration experience produces psychomotor sensitization, with similar characteristics to that produced by repeated, intermittent treatment with IP experimenter-administered drugs. It did. (1) Psychomotor sensitization was expressed both early (one day; WD1) and late (30 days, WD30) after the discontinuation of IntA self-administration, but was more robust at WD30 than WD1. Interestingly, following LgA self-administration experience psychomotor sensitization was only evident at WD30. (2) IntA self-administration experience produced greater psychomotor sensitization in female than male rats. (3) IntA self-administration experience produced cross psychomotor sensitization to another psychomotor stimulant drug, *d*-amphetamine. These findings have a number of implications for thinking about how cocaine may change brain and behavior in ways that can promote the transition to addiction, as discussed below.

### The effect of IntA vs LgA cocaine self-administration on the induction of psychomotor sensitization as a function of withdrawal period

#### IntA

In many early studies psychomotor stimulant drugs were administered IP or SC by an experimenter, repeatedly and intermittently, and in such studies sensitization was often only apparent, or expressed more strongly, after a period of withdrawal (Robinson and Camp 1987; Paulson et al. 1991; Paulson and Robinson 1995; Grimm et al. 2001). Similar effects were seen here following IntA cocaine self-administration experience. At WD1 psychomotor sensitization was manifest by locomotor hyperactivity (Fig. 3). However, by WD30 locomotor hyperactivity was no longer evident, because of the emergence of focused stereotyped behaviors (Fig. 4). Of course, the latter reflects a greater drug effect (Lyon and Robbins 1975; Segal 1975), and therefore, is evidence of robust psychomotor sensitization. These results are similar to those seen after ShA cocaine self-administration experience (see Introduction for references), and during IntA sessions, as reported by Samaha and colleagues (Allain et al. 2017; Allain and Samaha 2018; Algallal et al. 2019).

It is clear that IntA cocaine self-administration experience is capable of sensitizing brain circuitry that mediates the psychomotor activating effects of cocaine. Of course, the different psychomotor activating effects of psychostimulant drugs, including locomotor hyperactivity and stereotyped behaviors, are thought to be mediated largely by DA projections from the midbrain to both the dorsal and ventral striatum (Creese and Iversen 1975; Kelly et al. 1975; Pijnenburg et al. 1976; Vezina and Kim 1999; Vanderschuren and Kalivas 2000). The behavioral data suggest, therefore, that IntA would also produce DA sensitization. Indeed, following IntA cocaine self-administration experience a single self-administered infusion of cocaine produces a greater increase in DA in the core of the nucleus accumbens *in vivo*, than in control rats with more limited cocaine self-administration experience (Kawa et al. 2019b). Furthermore, IntA cocaine self-administration sensitizes electrically-evoked DA release measured in the accumbens *ex vivo* (Calipari et al. 2013) and sensitizes the ability of cocaine to inhibit DA uptake (Calipari et al. 2013, 2015). Thus, data to date show that IntA experience produces both psychomotor and DA sensitization, similar to the effects produced by intermittent, noncontingent injections of cocaine or amphetamine (for reviews see, Robinson and Becker 1986; Kalivas and Stewart 1991; Stewart and Badiani 1993; Vezina 2004).

#### LgA

Data on the ability of LgA or high dose cocaine self-administration experience to induce psychomotor sensitization are more mixed. When animals were tested soon after the discontinuation of LgA self-administration there are reports that the psychomotor activating effects of cocaine are actually attenuated (i.e., show tolerance), consistent with reports that the ability of cocaine to increase extracellular DA *in vivo* is also decreased, as is its ability to inhibit DA uptake (Calipari et al. 2013, 2014a). On the other hand, there are also reports of no tolerance to either cocaine’s psychomotor activating effects (Ahmed and Cador 2006), or effects on extracellular DA (Ahmed et al. 2004; Kawa et al. 2019b), following LgA experience, relative to ShA experience. When animals have been tested longer after the discontinuation of LgA experience (at least 14 days) studies are also mixed as to whether psychomotor sensitization is expressed. Some studies report that psychomotor sensitization is not evident, even after long periods of withdrawal (Ben-Shahar et al. 2004, 2005), and one study even reports evidence of tolerance to the psychomotor activating effects of cocaine at WD60 (Ben-Shahar et al. 2005). Others have reported that long after the discontinuation of LgA rats express similar sensitization to ShA rats (Knackstedt and Kalivas 2007), or even especially robust psychomotor sensitization (Ferrario et al. 2005), as was found here.

What might account for the very different effects of LgA experience, especially after a long period of withdrawal? One possibility is that in some studies animals were tested for psychomotor sensitization in a context where they had never experienced the drug. Under some circumstances the expression of psychomotor sensitization can be very context-specific (Badiani et al. 1995; Crombag et al. 2000), so this may account for some of the negative findings. Indeed, this may account for why in the current study males expressed sensitization within the self-administration chambers, but not when tested outside this environment (compare Figs. 7 and 8). Perhaps more important, however, is that some studies relied on a single measure of psychomotor activity, locomotion, and this can lead to spurious conclusions. As shown here, and by Ferrario et al (2005), at WD30 measures of locomotor activity did not reveal evidence of sensitization. But that was because of the emergence of focused stereotyped behaviors, which are indicative of a stronger (i.e., sensitized), not weaker, drug effect. Therefore, analysis of a range of behaviors (captured by the rating scale) and head movements (i.e., focused stereotypy) revealed that LgA experience had in fact produced very robust psychomotor sensitization. The dangers of relying solely on measures of locomotor activity in studies of psychomotor sensitization are discussed in detail by Ferrario et al. (2005) and Flagel & Robinson (2007), and they suggest that when only locomotor activity is assessed negative results have to be interpreted with great caution.

In summary, some, but not all, studies are consistent with the idea that LgA cocaine self-administration may attenuate the psychomotor activating effects of cocaine when animals are tested soon after the discontinuation of self-administration, but when tested after a period of withdrawal very robust psychomotor sensitization is manifest (Ferrario et al. 2005; present study). This is consistent with the effects of repeated treatment with high doses of psychostimulant drugs administered by an experimenter, when sensitization may not be expressed early after the discontinuation of drug treatment, but become evident after a few weeks of withdrawal (Robinson and Camp 1987; Paulson et al. 1991; Paulson and Robinson 1995; Grimm et al. 2001). One reason the psychomotor effects may change over time in this way may be because both tolerance- and sensitization-related neuroadaptations can be present at the same time (e.g., Izenwasser and French 2002), but tolerance-related adaptations can mask the expression of sensitization – sensitization is expressed only as tolerance-related adaptations subside. This interpretation is supported by an interesting study by Dalia et al. (1998). Briefly, rats previously expressing psychomotor sensitization were implanted with osmotic pumps that administered cocaine continuously for seven days. These rats expressed behavioral tolerance the day after removal of the pump, but by day 10 post-pump removal, the rats yet again expressed behavioral sensitization to cocaine, as tolerance waned.

### Sex differences in the effects of IntA experience on psychomotor sensitization

#### Self-Administration

Both male and female rats were initially trained to self-administer cocaine using an “infusion criteria” procedure. Under these conditions there was no sex difference in the acquisition of self-administration, consistent with a previous study using the same procedure (Kawa and Robinson 2019). It should be noted, however, that when tested under free access conditions females have been reported to more readily acquire cocaine self-administration (Lynch and Carroll 1999; Hu et al. 2004; although see Algallal et al. 2019). Here, males and females consumed the same amount of drug during acquisition, because each rat was allowed to take a predetermined and fixed number of injections. However, when tested using the ‘threshold’ procedure females consumed more cocaine than males at low doses (Fig. 5). This is consistent with reports that females are more motivated and/or consume more cocaine post-acquisition than males (Roberts et al. 1989; Cummings et al. 2011; Kawa and Robinson 2019). Therefore, in order to ensure that there were no group differences in cocaine consumption during IntA (which could impact the degree of sensitization) both males and females were limited to 3 cocaine injections during each Drug-Available period of the IntA-Limited self-administration procedure used in Exp 2.

#### Psychomotor sensitization

For these studies locomotor activity was monitored during IntA-Limited self-administration sessions and video was recorded throughout. Consistent with the absence of sex differences in cocaine consumption during IntA-Limited, there were no sex differences in the psychomotor activating effects of self-administered cocaine on the first day of IntA-Limited training. However, with increasing IntA-Limited experience a clear sex difference in the psychomotor activating effects of self-administered cocaine emerged, with females showing much more robust psychomotor sensitization than males (Fig 7; also see, Algallal et al. 2019). This sex difference was also evident on a probe test when rats with IntA-Limted experience were compared to a control group that had acquired cocaine self-administration and underwent the ‘threshold’ test, but did not have IntA-Limited experience (Fig. 8). Thus, both within-subjects and between-subjects comparisons revealed that following IntA-Limited females expressed greater psychomotor sensitization than males. These findings are consistent with many early studies reporting that experimenter-administered cocaine or amphetamine produces greater psychomotor sensitization in females than males (Glick and Hinds 1984; Robinson 1984; Van Haaren and Meyer 1991; for review, Becker et al. 2006). They are also consistent with a report that IntA experience produces more robust incentive-sensitization in females than males (Kawa and Robinson 2019). This greater propensity for sensitization in females may contribute to the more rapid emergence of problematic drug use (the ‘telescoping effect’) in women, described in clinical studies (Anglin et al. 1987; Kosten et al. 1993; Brady and Randall 1999).

### IntA experience and cross-sensitization

There are a number of early studies reporting psychomotor cross-sensitization between the two prototypical psychomotor stimulant drugs, amphetamine and cocaine, when they were administered by an experimenter (Shuster et al. 1977; Akimoto et al. 1990; Schenk et al. 1991; Hirabayashi et al. 1991; Bonate et al. 1997; Shanks et al. 2015). However, all the studies reporting that treatment with cocaine induces psychomotor sensitization to amphetamine involved the use of IP injections (Akimoto et al. 1990; Hirabayashi et al. 1991; Shanks et al. 2015). We first asked, therefore, whether experimenter-administered *IV* injections of cocaine would (1) induce psychomotor sensitization and (2) cross-sensitization to IV amphetamine. They did. We next asked whether IntA cocaine self-administration experience would also produce psychomotor cross-sensitization to IV amphetamine, and it did. This psychomotor cross-sensitization may be due to cross-sensitization of DA activity. IntA cocaine self-administration experience not only sensitizes the effects of cocaine at the DAT, but produce cross-sensitization to amphetamine’s actions at the DAT as well. Indeed, IntA (but not LgA) cocaine self-administration experience increases the ability of *amphetamine* to inhibit DA uptake (Calipari et al. 2014b). Interestingly, the noncontingent administration of amphetamine, which produces psychomotor sensitization, also facilitates escalation of intake when rats were later allowed to self-administer cocaine (Ferrario and Robinson 2007). The phenomenon of cross-sensitization may help explain why the use of one drug of abuse increases the probability that others will be abused as well, and why polydrug use is so common in people with substance abuse disorders (Schenk 2002).

### Conclusions and implications for theories of addiction

As mentioned in the Introduction, preclinical studies using LgA self-administration procedures have been cited in support of the idea that addiction is due, in part, to tolerance, leading to a drug-induced *hypo*dopaminergic state, and drug-seeking behavior is motivated to overcome this ‘DA deficiency’ (e.g., Koob and Volkow 2016; Volkow et al. 2016). Indeed, when tested soon after the discontinuation of LgA, or other high dose procedures, there may be tolerance to cocaine’s psychomotor activating effects, and its ability to increase DA neurotransmission (Ferris et al. 2011; Calipari et al. 2013, 2014a), although evidence for this is mixed (Ahmed et al. 2004; Kawa et al. 2019b and current results). However, after longer periods of withdrawal (30 days) robust psychomotor sensitization is evident (Ferrario et al. 2005; present study). We are not aware of any studies on the effects of LgA on DA neurotransmission after long periods of withdrawal. But, there are many studies showing that animals with prior LgA experience show enhanced glutamatergic transmission in the ventral striatum after long periods of withdrawal (Wolf 2016), when psychomotor sensitization is expressed, and when rats show especially robust reinstatement of drug-seeking behavior – so-called ‘incubation of craving’ (Grimm et al. 2001).

Most importantly, there is now considerable evidence that IntA cocaine self-administration, which is thought to better reflect human patterns of use, is more effective than LgA in producing addiction-like behavior, despite much lower levels of total drug consumption. IntA also produces robust psychomotor sensitization (present study), incentive-sensitization and DA sensitization (for reviews see, Allain et al. 2015; Kawa et al. 2019a), which is consistent with an incentive-sensitization view of addiction (Robinson and Berridge 1993). It will be interesting, therefore, to determine if IntA experience produces similar glutamatergic plasticity as LgA, and whether it is related to the expression of psychomotor sensitization and/or the robust reinstatement of drug-seeking that is evident even after short periods of withdrawal from IntA (although see Ferrario et al. 2010). Of course, it is also possible that the addiction-like behavior and psychomotor sensitization produced by IntA experience has a different neurobiological basis than that produced by LgA. If this is the case it would have important implications for thinking about how drugs may change brain and behavior in ways that promote a transition from casual drug use to the problematic patterns of use that define addiction.

## Acknowledgements

The authors thank Kendall Coden, Jack Hildenbrand, Joyce Xia and Lauren Longyear for their contributions to data collection and/or analysis. We thank NIDA for the gift of cocaine from the NIDA drug supply program. This research was supported by grant RO1 DA044204 from NIDA to TER and CRF and R21 DA045277 to CRF.

